# Surveillance of densoviruses and mesomycetozoans inhabiting grossly normal tissues of three New Zealand asteroid species

**DOI:** 10.1101/2020.10.08.331132

**Authors:** Ian Hewson, Mary A. Sewell

## Abstract

Asteroid wasting events and mass mortality have occurred for over a century. We currently lack a fundamental understanding of the microbial ecology of asteroid disease, with disease investigations hindered by sparse information about the microorganisms associated with grossly normal specimens. We surveilled viruses and protists associated with grossly normal specimens of three asteroid species (*Patiriella regularis, Stichaster australis, Coscinasterias muricata*) on the North Island, New Zealand, using metagenomes prepared from virus and ribosome-sized material. We discovered several densovirus-like genome fragments in our RNA and DNA metagenomic libraries. Subsequent survey of their prevalence within populations by quantitative PCR (qPCR) demonstrated their occurrence in only a few (13 %) specimens (n = 36). Survey of large and small subunit rRNAs in metagenomes revealed the presence of a mesomycete (most closely matching *Ichthyosporea* sp.). Survey of large subunit prevalence and load by qPCR revealed that it is widely detectable (80%) and present predominately in body wall tissues across all 3 species of asteroid. Our results raise interesting questions about the roles of these microbiome constituents in host ecology and pathogenesis under changing ocean conditions.

## INTRODUCTION

Recent and renewed interest in echinoderm microbiome ecology has revealed the paucity in understanding of the roles of the microbial community in host biology and ecology; particularly with respect to negative impacts such as mass mortality. Asteroid mass mortality due to a condition termed “sea star wasting disease” (also known as “asteroid idiopathic wasting syndrome”) has occurred in the northeast Pacific starting in 2013 [1], and in Port Phillip Bay, Australia and Shandong Province, China in 2014 [2]. Indeed, wasting has been observed for over a century [3]. Microbiological investigation of wasting asteroids initially indicated the presence of the Asteroid ambidensovirus 1 [4] (known at the time as Sea Star associated Densovirus or SSaDV; [1]), and wasted asteroids were inhabited by a suite of cultivable copiotrophic (i.e. bacteria that rapidly consume abundant organic matter) bacteria [5, 6]. Firm microbial associations with sea star wasting remain elusive, similar to other echinoderm diseases (reviewed in [7]). Despite the lack of conclusive disease etiology, previous work has highlighted distinct microbiome associations with echinoderms [8], building on previous microscopic and cultivation-based studies [9–11]. These surveys suggest that echinoderms may harbor an underexplored diversity of microorganisms. Environmental perturbation under future climate scenarios may shift the relationship between these microorganisms and their hosts [12]. Hence, there is value in surveying the diversity and prevalence of microorganisms associated with grossly normal specimens, which may then inform future marine disease event investigations, when and if they occur.

A grand challenge in surveying microbial eukaryotic microorganisms associated with metazoa using PCR-based approaches is that well-conserved marker genes (e.g. ribosomal RNAs) are shared between symbiotic partners. Hence, unbiased surveys of host-associated protists using PCR amplification-based approaches are limited. Modified primer design to exclude metazoan partners, using primers distinct to expected taxonomic groups (e.g. fungal ITS; [13]), and the use of blocking PCR primers [14, 15] may alleviate this burden, but demand *a priori* knowledge of native protistan diversity. The study of viral diversity associated with metazoan hosts has been approached by two methods. First, viral genomes have been recovered from deeply sequenced host transcriptomes [16, 17]. This approach provides key information about expressed host genes in addition to a wealth of viral diversity, including deeply-branching viral genotypes across a wide range of invertebrate hosts [17]. A second approach enriches for viruses by physical size and capsid-induced protection from nucleases [18]. Here, viral metagenomes are typically prepared using a homogenization-size exclusion-nuclease approach, where tissues are normally ‘cleaned’ (washed) of putative epibionts [19]. Viral metagenomes prepared using this approach have potential to yield more information than viruses alone, since only a tiny fraction (typically < 5%) of metavirome sequence space is annotated as viruses [20] and the remaining sequence space is believed to mostly reflect host RNAs. Ribosomes, which are typically 25 – 30 nm in diameter, are also liberated from cells during homogenization, pass through the filters typically used in metavirome preparation, and transcript RNAs may be protected from nucleases used to digest co-extracted nucleic acids. Thus, ribosomal RNAs are well represented in viral metagenomes and may include protistan, bacterial and archaeal components of the host-associated microbiome. Comparison of non-viral sequences in viral metagenomes against rRNA databases can be used to study microbiome constituents that are inaccessible or impractically studied by PCR-based approaches.

The goal of the present study was to identify viruses and protists in common New Zealand asteroids by surveying virus- and ribosome-sized RNAs, and use this information to guide survey of microbial prevalence within and between populations and between tissue types. We discovered several densovirus genome fragments in two species of asteroid, but these were only detected at low prevalence within the populations studied by quantitative PCR. We also discovered fungal, mycetozoan and mesomycetozoan constituents of the asteroid microbiome. A mesomycetozoan similar to a fish pathogen was prevalent in all asteroids tested, and bore highest loads in body wall samples, suggesting it may be a common constituent of the asteroid microbiome.

## MATERIALS AND METHODS

### Sample collection

Asteroid samples (n = 77 individuals across 3 species) were collected for metagenomic investigation of viral diversity and viral prevalence at several locations on the North Island, New Zealand, in January and February 2018 (Table 1). Asteroids were collected by hand, either from the intertidal or subtidal, and immediately placed into individual plastic bags, which were transported to the laboratory for dissection in a cooler (Fig. 1). The taxonomic identity and arm length of individuals was recorded for each specimen. Coelomic fluid was withdrawn from individuals using a 5 mL syringe fitted with a sterile 25G needle. Body wall tissues were removed by sterile (5 mm) biopsy punch. Gonads and pyloric caeca were dissected from coelomic cavities by first creating an incision into the coelomic cavity using clean disposable razor blades, then using sterilized forceps to remove small (∼ 2 – 4 mm) sections of these tissues. All tissue and coelomic fluid samples were preserved in RNALater at a ratio of 2:1 (vol:vol), refrigerated, and transported to the laboratory at Cornell University for further processing, which occurred within 4 months of collection.

**Table 1:** Sampling locations, species and morphological characteristics of asteroids collected as part of this study. Samples collected at Ti Point were collected subtidally by SCUBA Diver, while those collected elsewhere were collected intertidally. RL = Ray length, SE = Standard Error.

| Location | Latitude | Longitude | Date | Species | n | RL (cm) | RL SE |
| --- | --- | --- | --- | --- | --- | --- | --- |
| Piha, Auckland | 36.9597 S | 174.4628 E | 1/22/2018 | *Stichaster australis* | 19 | 13.58 | 0.56 |
| Ti Point, Northland | 36.3178 S | 174.6178 E | 1/27/2018 | *Coscinasterias muricata* | 17 | 7.42 | 1.10 |
| Matheson's Bay, Northland | 36.3011 S | 171.8011 E | 1/27/2018 | *Patiriella regularis* | 20 | 2.81 | 0.17 |
|  |  |  |  | *Coscinasterias muricata* | 1 | 10.00 |  |
| Scorcher Bay,<br>Wellington | 41.3078 S | 174.8325 E | 2/15/2018 | *Patiriella regularis* | 20 | 1.89 | 0.11 |

**Figure 1:**
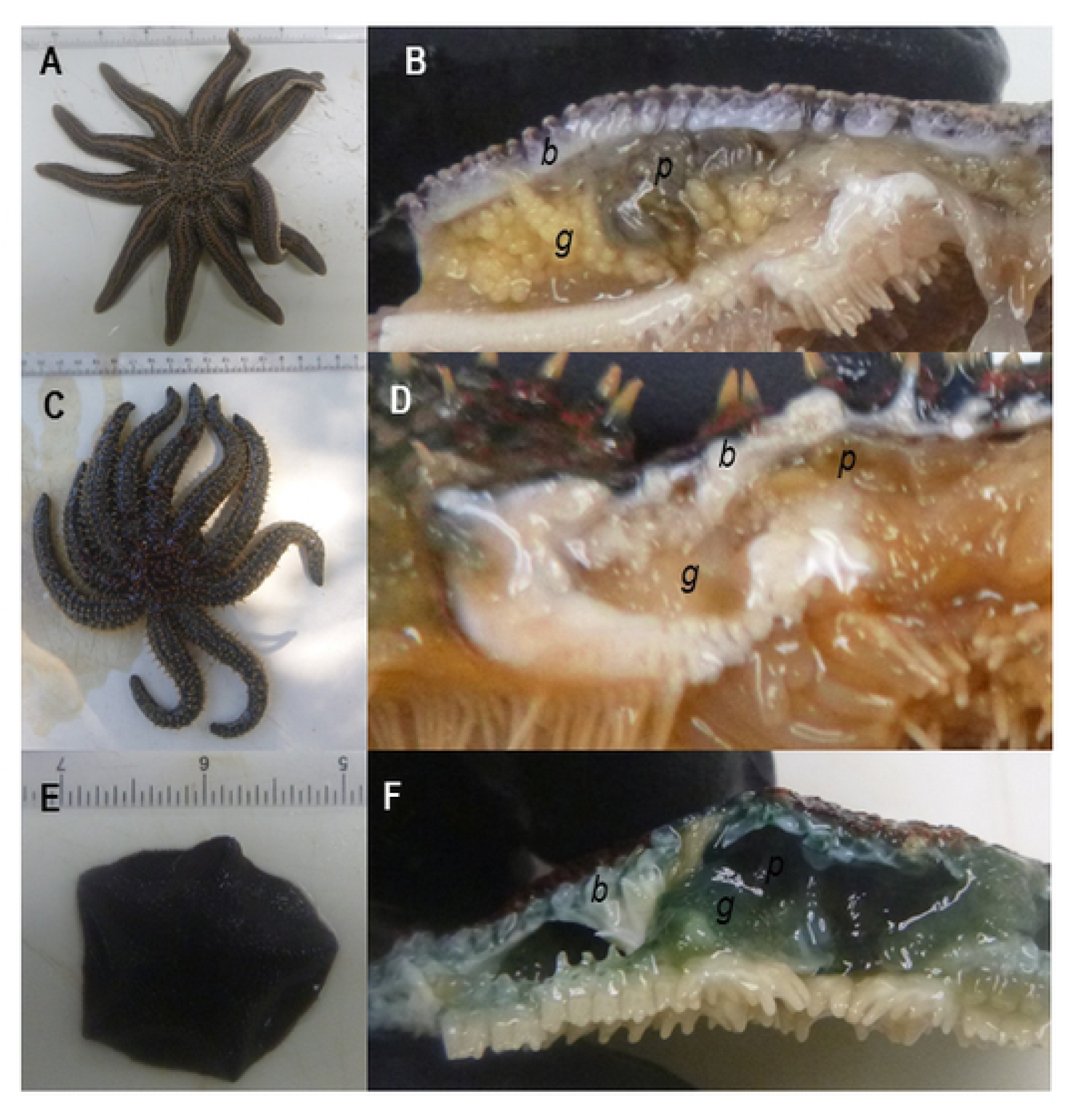
Sampled specimens of *Stichaster australis* (A-B), *Coscinasterias muricata* (C-D) and *Patiriella regularis* (E-F). Viral metagenomes were prepared from body wall (*b*) samples collected by biopsy punch. Additional specimens of gonad (*g*) and pyloric caeca (*p*) were collected for quantification of viral genotypes and the mesomycetozoan.

### Metavirome Preparation

Three body wall biopsy samples from each species were selected for viral metagenomics (one each from *Stichaster australis, Coscinasterias muricata* and *Patiriella regularis*; Table 2). For each sample, the biopsy punch was removed from RNALater and subject to the workflow detailed in [19] with modifications by Ng et al [21] and Hewson et al. [22].

**Table 2:** Library characteristics prepared from asteroids in Northland and Auckland region, January 2018.

| Species | Date | Total Reads | Assembled Reads | Total Contigs | Viral Contigs |
| --- | --- | --- | --- | --- | --- |
| *Coscinasterias muricata* | 1/27/2018 | 3,867,602 | 981,140 | 27,032 | 2 |
| *Stichaster australis* | 1/22/2018 | 1,673,102 | 681,086 | 2,170 | 0 |
| *Patiriella regularis* | 1/27/2018 | 1,635,372 | 301,574 | 6,332 | 2 |

**Table 3:**
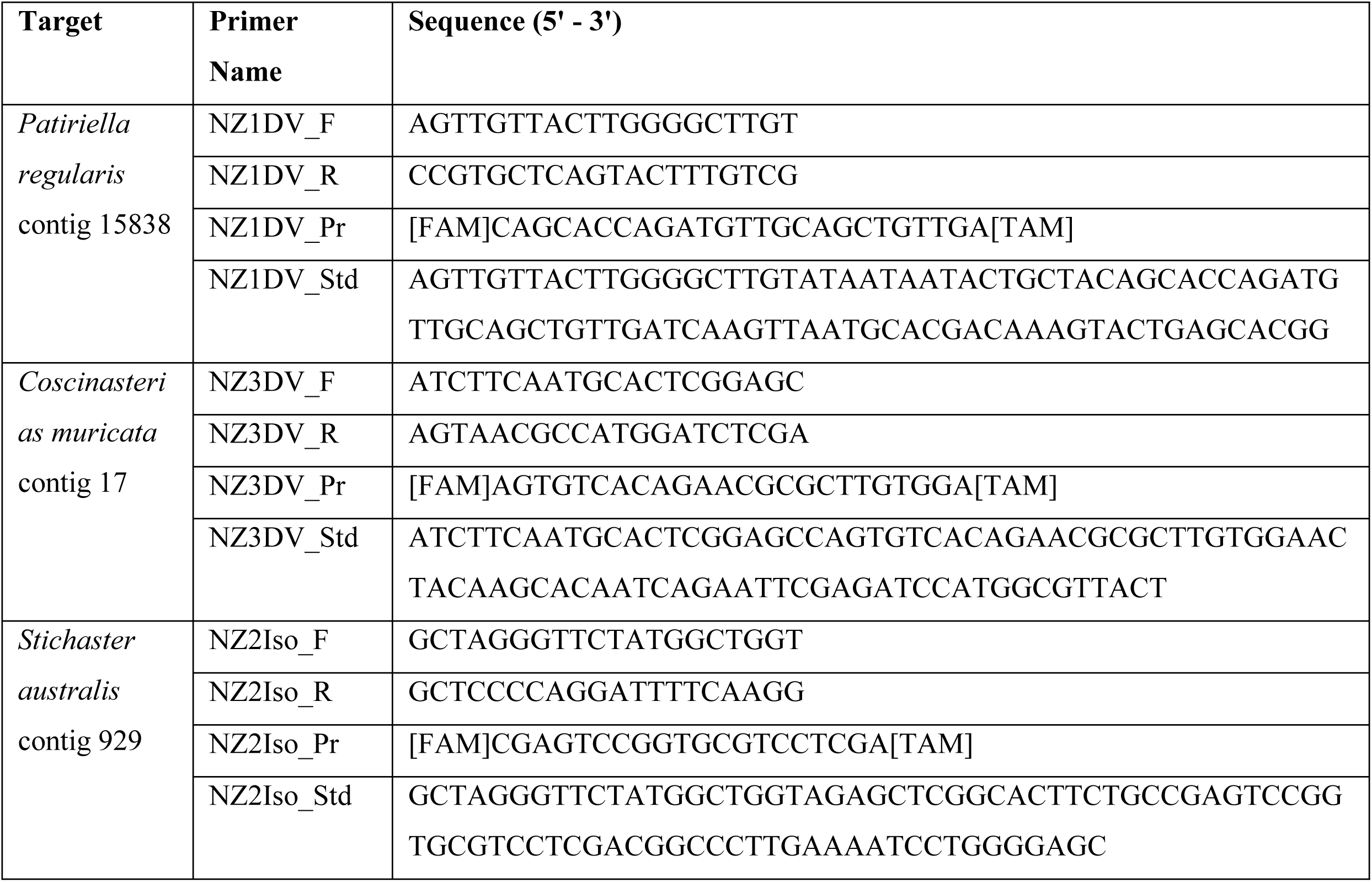
Primers and probes used in this study to examine the prevalence and load of densovirus and mesomycetozoan-like contiguous sequences.

Briefly, the sample was homogenized by bead beating (Zymo Bead Beater tubes) in 1 mL of 0.02 µm-filtered PBS. The sample was filtered through a 0.2 µm PES syringe filter. The filtrate was treated with DNAse I (5 U; Thermo Fisher Scientific), RNAse One (50 U; Promega) and Benzonase (250 U; Sigma-Aldrich) for 3 h at 37^°^C in an attempt to remove co-extracted host nucleic acids. Enzyme activity was halted by treatment with 50 µM EDTA. RNA was extracted from the resulting purified viral fraction using the Zymo Mini RNA isolation kit, and subsequently amplified using the TransPlex WTA2 (Sigma Aldrich) kit. We did not standardize template quantity of extracted RNA (2 µl) prior to amplification. Amplicons were quantified using PicoGreen and submitted for sequencing on a Illumina MiSeq (2 × 250 bp paired-end) platform after TruSeq PCR-free library preparation at the Cornell Biotechnology Resource Center. Sequences have been deposited in the NCBI under BioProject PRJNA636826.

### Bioinformatic processing

Sequence libraries were initially trimmed for adapters and quality (ambiguous bases <2) using the CLC Genomics Workbench 4.0. Each of the 3 metaviromes were assembled separately using the CLC Genomic Workbench 4.0 native algorithm using a minimum overlap of 0.5 and similarity of 0.8. The resulting contig spectra was aligned against several boutique databases of RNA viruses as described elsewhere [22]. Because RNA viral metagenomes also capture ssDNA viruses [23], we also searched contig spectra by tBLASTx against a boutique database of densoviral genomes (complete genomes from NCBI using keyword “densovirus”). Sequence matches against any of these databases at an E-value <10^−20^ were further aligned against the non-redundant (nr) library at NCBI by BLASTx, and contigs discarded if they matched known bacterial or eukaryl proteins at a higher percentage and E-value than viruses. Uncertain amplification biases and variation in template RNA quantity preclude quantitative interpretation of metagenome constituents. Hence, analyses of metagenomes focused on detection of constituents and subsequent quantitative PCR of selected contigs.

### Quantitative PCR (qPCR) of densovirus genome fragments

TaqMan Primer/Probe sets were designed around two contiguous sequences matching the nonstructural proteins of densoviruses (*Coscinasterias muricata* contig 17 and *Patiriella regularis* contig 15838) and validated against an oligonucleotide standard (Table 2). DNA was extracted from 36 biopsy punch body wall samples (10 *Stichaster australis*, 3 *Coscinasterias muricata*, 13 *Patiriella regularis* from near Auckland and 10 *Patiriella regularis* from Scorcher Bay, Wellington) using the Zymo Tissue & Insect Kit. DNA was then subject to quantitative PCR (qPCR) in an Applied Biosystems StepOne Real-Time PCR machine. Each qPCR reaction comprised 1 X SSO Probes SuperMix (BioRad), and 200 pmol of each primer and probe (Table 2). Reactions were subject to a 10 minute incubation step at 50°C, followed by a 3 minute denaturation step at 94°C. Following hot start activation, reactions were subject to 50 cycles of heating to 94°C and annealing at 58°C, where fluorescence was measured at the conclusion of each thermal cycle. Reactions were run in duplicate against an 8-fold dilution (covering 10 to 10^8^ copies reaction^−1^). A positive detection of the virus was considered when both duplicates were within 1 Ct, and were considered “detected but not quantifiable” (DNQ) when one replicate generated a positive Ct but the other replicate failed to yield an amplicon.

### Investigation of eukaryote 18S and 28S rRNAs in metaviromes

Contiguous sequences generated from viral metagenomes (described above) were queried against the Silva database (version r132) of 16S/18S and 23S/28S rRNAs [24] by BLASTn and contigs matching at E<10^−10^ to 18S or 28S rRNAs were considered for further analysis. Matches meeting this criterion were then queried against the non-redundant database at NCBI. Matches to asteroid 18S and 28S rRNAs were removed from further consideration, as were matches to other metazoan rRNAs. The resulting contig spectra were aligned against close matches from NCBI using the CLC Sequence Viewer 8.0 (Qiagen).

### Investigation of mesomycetozoan tissue and species specificity

Quantitative PCR (qPCR) primers were designed around the 28S rRNA sequence matching *Ichthyosporea* sp. (*Stichaster* contig 929) and used to amplify body wall DNA extracts from 20 *Stichaster australis*, 6 *Coscinasterias muricata*, and 10 *Patiriella regularis*. Additionally, for each of the 20 *Stichaster australis*, samples of pyloric caeca and gonad were also examined for the presence and abundance of this sequence.

## RESULTS AND DISCUSSION

### Viruses associated with Asteroid Tissues

Metaviromes prepared from asteroid body wall samples contained between 1.6 – 3.9 million paired-end reads (Table 2). Assembly of these resulted in 2,170 to 27,032 contigs, where contigs recruited 18 – 41% of total reads. No RNA viruses were detected by alignment. However, alignment against densoviral genomes resulted in 2 contigs matching to the nonstructural gene 1 (NS1) and 2 contigs matching structural (VP) genes at E < 10^−15^ (Fig. 2). Three of these contigs – two from *Coscinasterias muricata* and one from *Patiriella regularis*-overlapped with ambidensovirus peptide sequences recovered from species of starfish collected worldwide. A further contig from *Patiriella regularis* matched a decapod penstyldensoviruses. Phylogenetic analyses based on NS1 revealed that *Coscinasterias muricata* contig 16413 was most similar to densoviruses recovered from *Asterias rubens* in Scotland [25] (Fig. 3). Phylogenetic analyses based on structural genes of the remaining viral contigs (*Coscinasterias muricata* contig 15838 and *Patiriella regularis* contig 3718) suggested that these were most similar to ambidensoviruses from molluscs [26, 27], insects [28], a crustacean [29], and human spinal fluid [30]. Quantitative PCR (qPCR) of the densovirus-like *Coscinasterias muricata* contig 15838 yielded only 3 DNQ results; two in *Coscinasterias muricata* (of 3 total surveyed) from Ti Point; and one *Stichaster australis* from Piha. qRT-PCR of *Patiriella* contig 17 yielded two DNQ results, both from *Patiriella regularis* collected at Matheson’s Bay. In no sample did we consistently detect the presence of either contig between replicate amplifications. This may be interpreted as indicating their very low copy number (<10) in DNA extracts.

**Figure 2:**
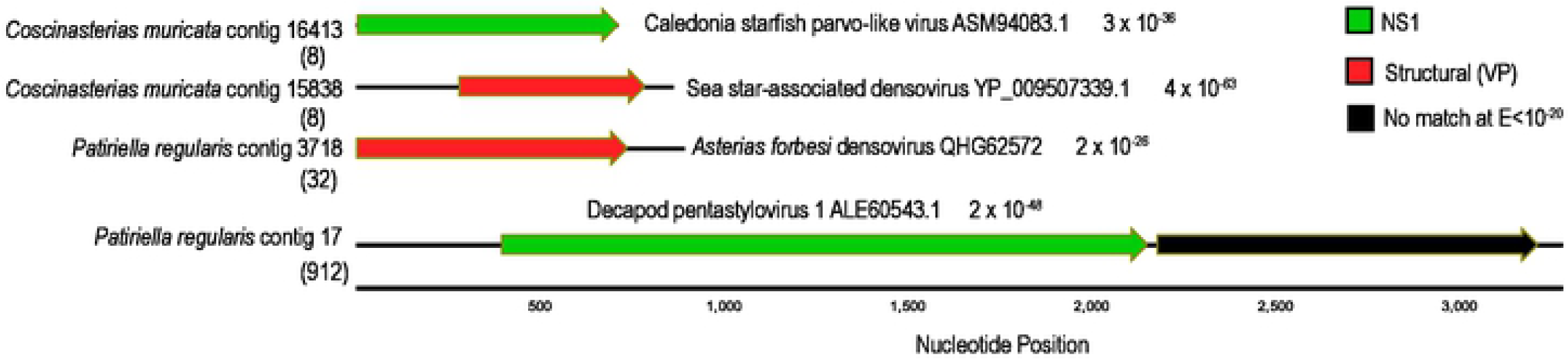
Contig map of densovirus-like genome fragments recovered from *Coscinasterias muricata* and *Patiriella regularis* viral metagenomes. The colors of arrows indicate densoviral gene, and the best match (by BLASTx against the non-redundant database at NCBI) along with e-value is indicated adjacent to each ORF. The black lines running through ORFs indicate total contig length. The numbers in brackets below each contig are the number of reads recruiting to the contig from the origin library.

**Figure 3:**
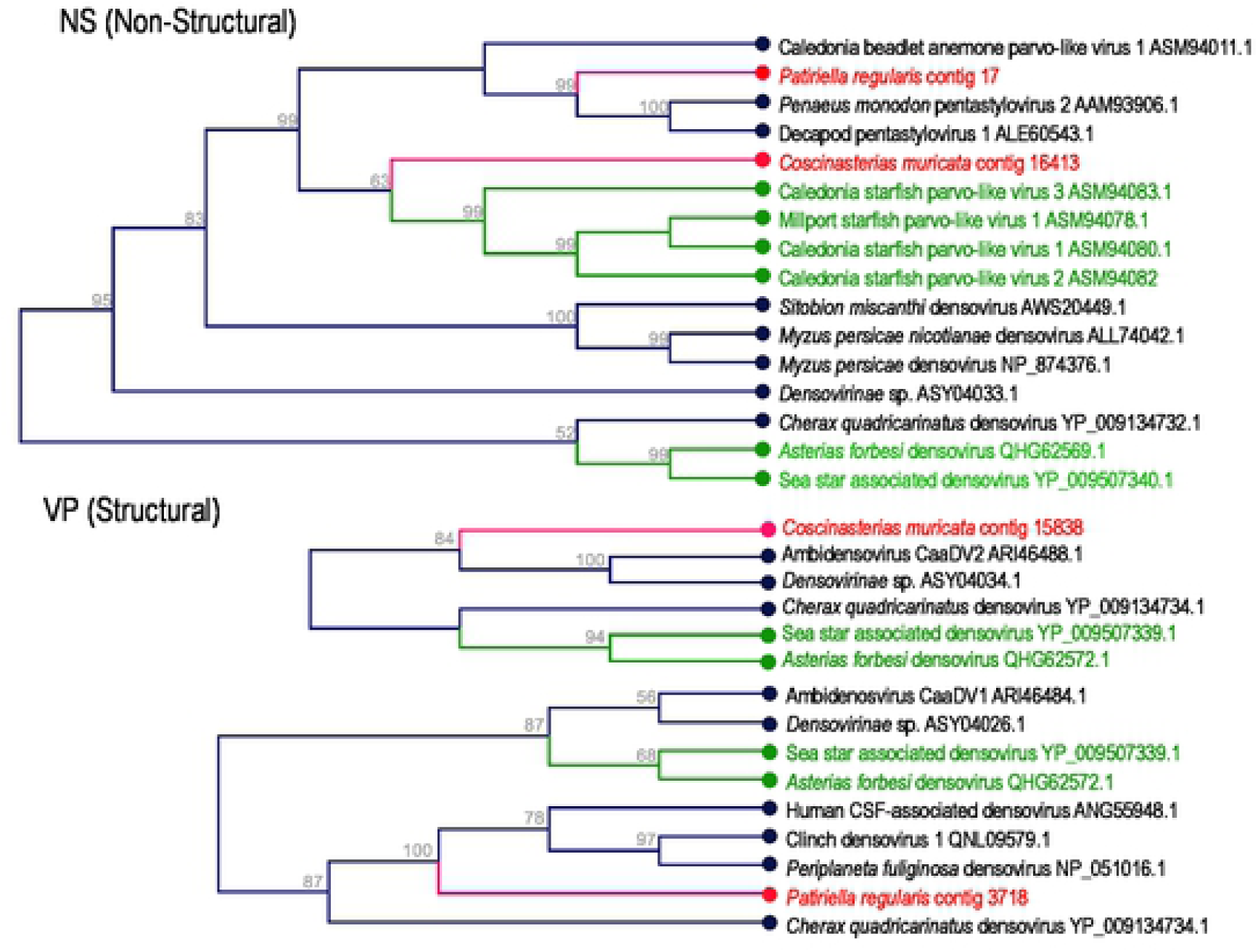
Phylogenetic representation of *Patiriella regularis* and *Coscinasterias muricata*-associated densoviral genome fragments. The trees are based upon 170 amino acid (Non-Structural), 211 amino acid (Structural; middle), and 104 amino acid (Structural; bottom) alignments performed using the CLC Sequence Viewer version 8.0. The trees were constructed with neighbor joining and Jukes-Cantor distance, where bootstrap values (1000 reps) are indicated above nodes. Red labels indicate sequences obtained in this study, while green labels indicate sequences obtained from asteroids in other studies.

The observation of densoviruses in these species was not surprising, since their recovery in other asteroids [1, 5, 23, 31] and urchins [32] suggests they may be a common constituent of echinoderm microbiomes. Parvoviruses form persistent infections in hosts [33, 34], and are highly prevalent and persistent in asteroid populations [31]. They are also widely endogenized in host genomes [35]. None of the densovirus-like contigs discovered in this survey represented complete genomes, so it is possible that these also represent endogenized densoviruses. However, flanking regions to their open reading frames did not match asteroid genomes, suggesting they were unlikely to be endogenized genome elements.

The pathology of densoviruses and significance in wasting diseases or other conditions is unclear. The copy number of Asteroid ambidensovirus-1 (SSaDV) and related densoviruses is elevated in wasting-affected *Pycnopodia helianthoides* [5]. However, histopathology [36] and other investigations [6, 23, 31] have failed to clinically connect densoviruses (or viruses in general) to sea star wasting disease. Densoviruses, like all parvoviruses, replicate in somatic cells. Infection in arthropods leads to respiratory impairment [37] and triggering of apoptosis [38], and has been linked to elevated mortality in crustacea [29, 39]. The discovery of a penstyldensovirus genome fragment in *Patiriella regularis* raises interesting questions about its role in host ecology. In penaeid shrimp, persistent infection by penstyldensoviruses delays mortality from white spot syndrome virus [40], suggesting densoviruses in general may play both detrimental and beneficial roles in host ecology. None of the asteroids sampled in this survey were grossly abnormal, and the low prevalence of the *Patiriella regularis* penstyldensovirus genome fragment in asteroid populations at our collection sites may indicate that these infections represent sub-clinical, or perhaps persistent infections which are unrelated to wasting or mass mortality.

### Protists associated with asteroids

A total of 15 contigs matched 18S and 28S rRNAs based on alignment. Of these, nine were fungal (five were Ascomycetes, four were Basidiomycetes), two were mycetozoan and one was mesomycetozoan (Fig. 4; Figs. S1-S4). The mesomycetozoan contiguous sequence (*Stichaster* contig 929) was most similar to a fish pathogen *Ichthyosporea* sp. ex *Tenebrio molitor* (Fig. 4). The abundance of this contiguous sequence was significantly higher in the body wall of *Stichaster australis* than in either *Coscinasterias muricata* (p = 0.019, Student’s t-test, df=4) or *Patiriella regularis* (p = 0.018, Student’s t-test, df=4) (Fig. 5). The abundance in epidermal tissues was also significantly higher in *Stichaster australis* than in either gonads or pyloric caeca (p = 0.013 and p=0.006, respectively, Student’s t-test, df=8). The contiguous sequence was detected in any quantity in 80 – 85% of all samples tested with no pattern with tissue specificity or species.

**Fig. 4:**
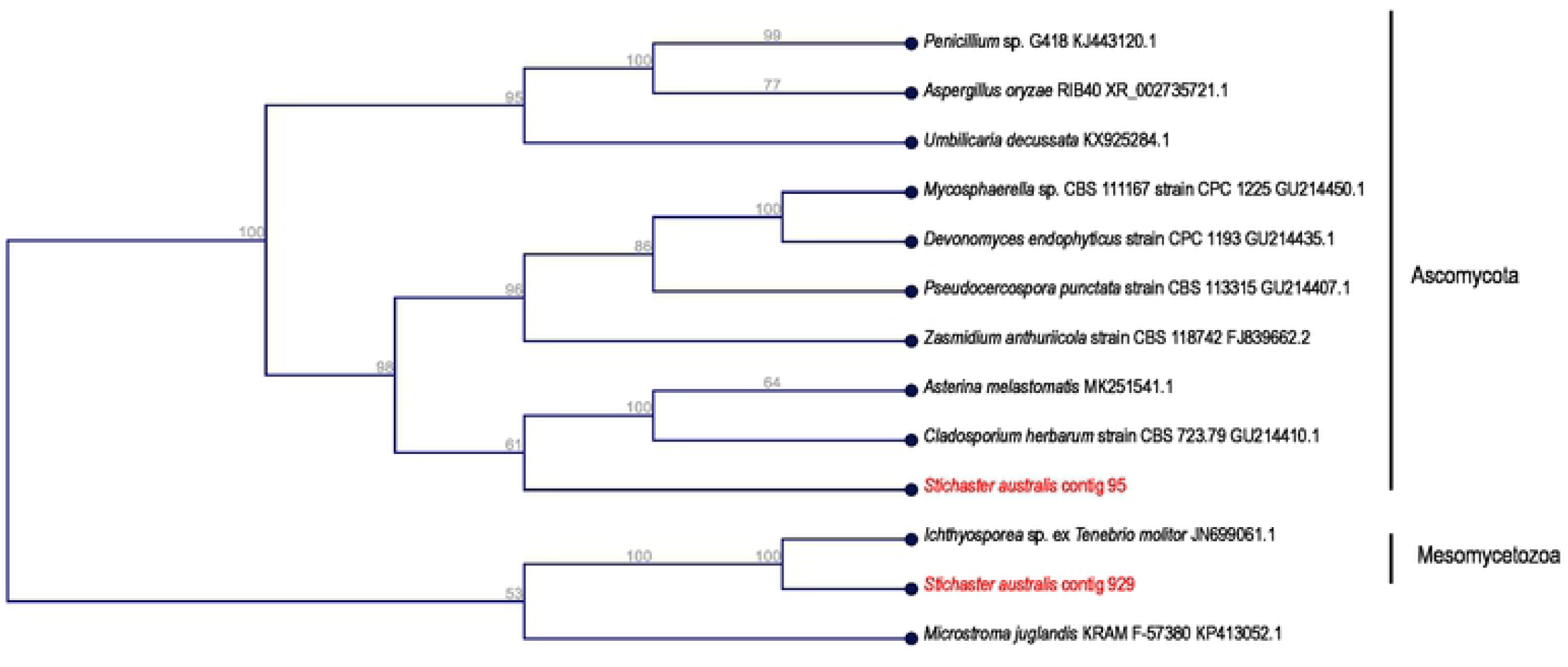
Phylogenetic representation of asteroid-associated 28S rRNA sequences in purified virus metagenomes. The tree was constructed by neighbor joining and based on an 849 nucleotide alignment of eukaryotic 28S rRNA. Shown are close matches by BLAST against the non-redundant database.

**Fig. 5:**
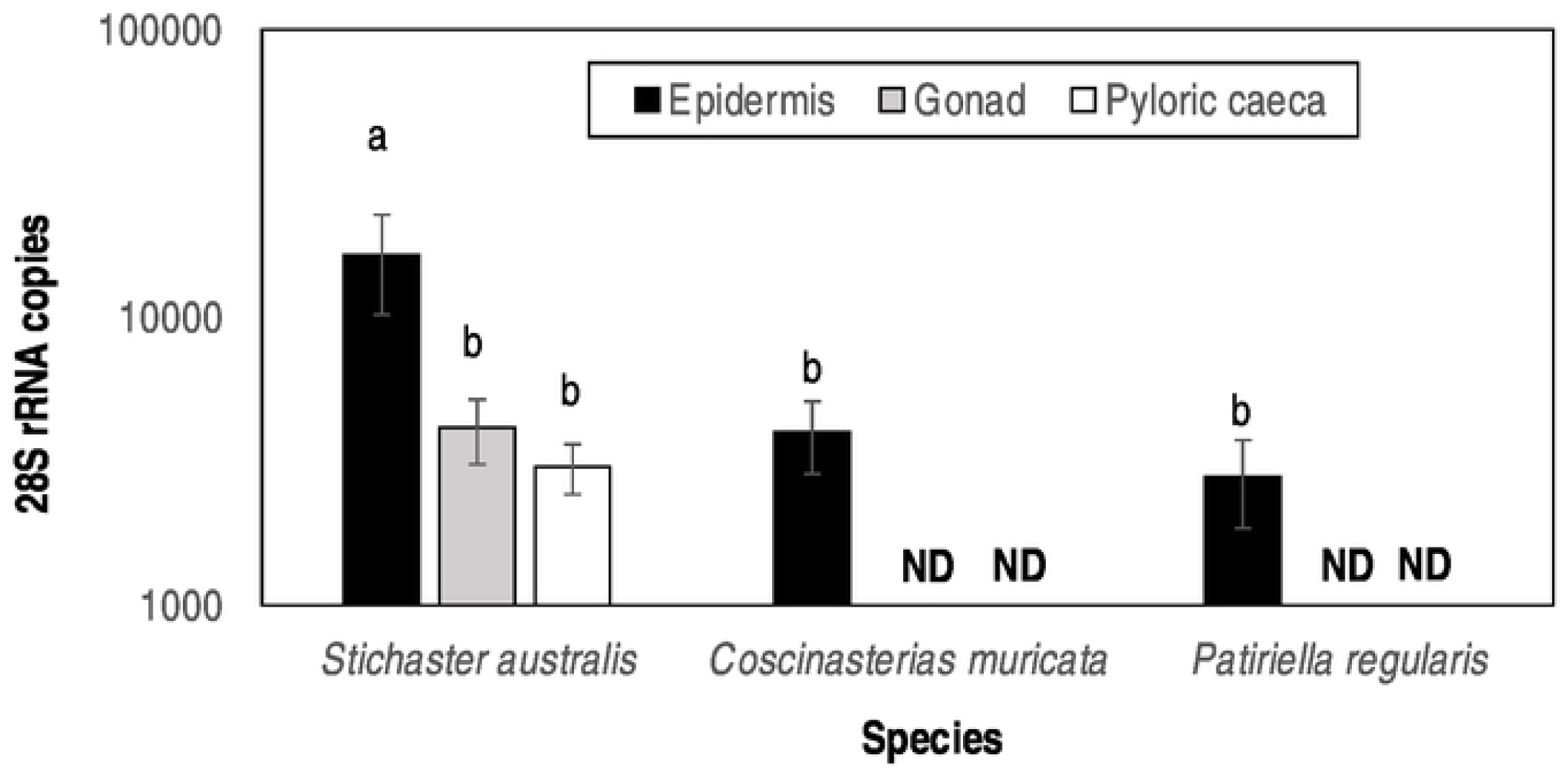
Mesomycetozoan 28S rRNA copies as determined by qPCR in asteroid tissues. a,b denotes significant difference (p < 0.025, Student’s t-test with Bonferroni correction for 2 comparisons).

The association of microbial eukaryotes, especially fungi and fungi-like protists, with echinoderms is not extensively documented in previous surveys. Hewson et al [22] reported the detection of totiviruses, which are fungal viruses, in several Holothuroidea. Similarly, Nerva et al [41] reported the mycovirome of fungi isolated from *Holothuria polii*. These reports suggest that fungi may be common constitutents of the sea cucumber microbiome. Wei et al [42] reported the cultivation of a symbiotic fungi most similar to *Penicillium* from an asteroid in China. Labyrinthulids have also been cultivated from the surface of wasting asteroids in the northeast Pacific [43]. However, their role in wasting pathology is unknown. There has been a body of work examining anti-fungal properties of asteroid extracts [44–47], suggesting that fungi discovered in this survey are unlikely to represent pathogens, but rather may be adapted to the chemical environment of their host. Further investigation of their roles in host chemical defense and dysbiosis is therefore warranted.

Mesomycetozoa are of interest since they represent the closest unicellular ancestor to multicellular animals [48]. They represent parasites of vertebrates [49–51] of which several, including *Ichthyosporea* spp. are aquatic. Aquatic mesomycetozoans infect fish and amphibians [50–53] and cause dermal disease. Mesomycetozoa may also form symbioses with their hosts (e.g. the mealworm *Tenebrio molitor;* [54] and other taxa [55, 56] (reviewed in [57]). Our observation of an *Icthyosporea*-like rRNA in *Stichaster* is the first report of this group in Asteroidea. The observations of greater load in epidermal tissues than internal organs suggests they may also form dermal infections, and their widespread occurrence in asteroid populations from the North Island of New Zealand suggests that mesomycetozoans are non-specific and broadly prevelant. Because we did not observe gross disease signs in any specimen, it is unlikely that this microorganism is a pathogen, but rather, they may represent a normal constituent of the host microbiome.

## CONCLUSIONS

To the best of our knowledge, this is the first investigation of viruses and mesomycetozoa associated with asteroids in New Zealand. Discovery of these taxa suggests an undiscovered bank of potential parasites or symbionts inhabiting echinoderms, and demands further investigation into their ecological roles. While we did not observe gross signs of disease in any specimen, we speculate that they may cause sub-clinical disease, or may interact with changing ocean conditions and give rise to more extensive disease events in the future. Our work demonstrates the value in unbiased surveys of microbiome constituents (i.e. microbial surveillance) which may inform future disease investigations by providing a picture of grossly normal microbiome constituents.

## ACKNOWLEDGEMENTS

The authors thank Richard Taylor for sample collection at Ti Point, and Christopher DeRito, Elliot Jackson and Kalia Bistolas for assistance with laboratory analyses. This work was supported by the United States National Science Foundation grants OCE-1737127 and OCE-1537111 to IH. The datasets generated during the current study are available in the NCBI repository under BioProject PRJNA636826.

## SUPPORTING INFORMATION CAPTIONS

**Fig. S1:** Phylogenetic representation of asteroid-associated 18S rRNA sequences in purified virus metagenomes. The tree was constructed by neighbor joining and based on an 689 nucleotide alignment of eukaryotic 18S rRNAs. Shown are close matches by BLAST against the non-redundant database.

**Fig. S2:** Phylogenetic representation of asteroid-associated ascomycete 28S rRNA sequences in purified virus metagenomes. The tree was constructed by neighbor joining and based on an 368 nucleotide alignment of eukaryotic 28S rRNAs. Shown are close matches by BLAST against the non-redundant database.

**Fig. S3:** Phylogenetic representation of asteroid-associated 28S rRNA sequences in purified virus metagenomes. The tree was constructed by neighbor joining and based on a 481 nucleotide alignment of eukaryotic 28S rRNAs. Shown are close matches by BLAST against the non-redundant database.

**Fig. S4:** Phylogenetic representation of asteroid-associated ascomycete 28S rRNA sequences in purified virus metagenomes. The tree was constructed by neighbor joining and based on a 506 nucleotide alignment of eukaryotic 28S rRNAs. Shown are close matches by BLAST against the non-redundant database.

## REFERENCES

1. Hewson I, Button JB, Gudenkauf BM, Miner B, Newton AL, Gaydos JK, et al. Densovirus associated with sea-star wasting disease and mass mortality. Proc Nat Acad Sci USA. 2014;111:17276–83.

2. Hewson I, Sullivan B, Jackson EW, Xu Q, Long H, Lin C, et al. Perspective: Something old, something new? Review of wasting and other mortality in Asteroidea (Echinodermata). Front Mar Sci. 2019;6(406). doi: 10.3389/fmars.2019.00406.

3. Mead AD. Twenty-eighth annual report of the commissioners of inland fisheries, made to the General Assembly at its January session, 1898. Providence: 1898.

4. Walker PJ, Siddell SG, Lefkowitz EJ, Mushegian AR, Dempsey DM, Dutilh BE, et al. Changes to virus taxonomy and the International Code of Virus Classification and Nomenclature ratified by the International Committee on Taxonomy of Viruses (2019). Archiv Virol. 2019;164(9):2417–29. doi: 10.1007/s00705-019-04306-w.

5. Hewson I, Bistolas KSI, Quijano Carde EM, Button JB, Foster PJ, Flanzenbaum JM, et al. Investigating the complex association between viral ecology, environment and Northeast Pacific Sea Star Wasting. Front Mar Sci. 2018; https://doi.org/10.3389/fmars.2018.00077.

6. Aquino CA, Besemer RM, DeRito CM, Kocian J, Porter IR, Raimondi PT, et al. Evidence that non-pathogenic microorganisms drive sea star wasting disease through boundary layer oxygen diffusion limitation. bioRxiv. 2020:doi: https://doi.org/10.1101/2020.07.31.231365

7. Hewson I. Technical pitfalls that bias comparative microbial community analyses of aquatic disease. Dis Aquat Organ. 2019;137(2):109–24. doi: 10.3354/dao03432.

8. Jackson EW, Pepe-Ranney C, Debenport SJ, Buckley DH, Hewson I. The microbial landscape of sea stars and the anatomical and interspecies variability of their microbiome. Front Microbiol. 2018;9:12. doi: 10.3389/fmicb.2018.01829.

9. Kelly MS, Barker MF, McKenzie JD, Powell J. The incidence and morphology of subcuticular bacteria in the echinoderm fauna of New Zealand. Biol Bull. 1995;189(2):91–105. doi: 10.2307/1542459.

10. Kelly MS, McKenzie JD. Survey of the occurrence and morphology of sub-cuticular bacteria in shelf echinoderms from the northeast Atlantic Ocean. Mar Biol. 1995;123(4):741–56. doi: 10.1007/bf00349117.

11. Holland ND, Nealson KH. Fine structure of the echinoderm cuticle and the sub-cuticular bacteria of echinoderms. Acta Zool. 1978;59(3-4):169–85. doi: 10.1111/j.1463-6395.1978.tb01032.x.

12. Egan S, Gardiner M. Microbial dysbiosis: Rethinking disease in marine ecosystems. Front Microbiol. 2016;7. doi: 10.3339/fmicb.2016.00997.

13. Nuñez-Pons L, Work TM, Angulo-Preckler C, Moles J, Avila C. Exploring the pathology of an epidermal disease affecting a circum-Antarctic sea star. Sci Rep. 2018;8:12. doi: 10.1038/s41598-018-29684-0.

14. Leray M, Agudelo N, Mills SC, Meyer CP. Effectiveness of annealing blocking primers versus restriction enzymes for characterization of generalist diets: unexpected prey revealed in the gut contents of two coral reef fish species. PLoS One. 2013;8(4):e58076–e. doi: 10.1371/journal.pone.0058076.

15. Vestheim H, Jarman SN. Blocking primers to enhance PCR amplification of rare sequences in mixed samples - a case study on prey DNA in Antarctic krill stomachs. Front Zool. 2008;5:12-. doi: 10.1186/1742-9994-5-12.

16. Holmes EC. The expanding virosphere. Cell Host Microbe. 2016;20(3):279-80.

17. Shi M, Lin X-D, Tian J-H, Chen L-J, Chen X, Li C-X, et al. Redefining the invertebrate RNA virosphere. Nature. 2016;540:539–43.

18. Ng TFF, Manire C, Borrowman K, Langer T, Ehrhart L, Breitbart M. Discovery of a novel single-stranded DNA virus from a sea turtle fibropapilloma by using viral metagenomics. J Virol. 2009;83(6):2500–9. doi: Doi 10.1128/Jvi.01946-08.

19. Thurber RV, Haynes M, Breitbart M, Wegley L, Rohwer F. Laboratory procedures to generate viral metagenomes. Nat Protocol. 2009;4(4):470–83. doi: DOI 10.1038/nprot.2009.10.

20. Correa AMS, Welsh RM, Thurber RLV. Unique nucleocytoplasmic dsDNA and +ssRNA viruses are associated with the dinoflagellate endosymbionts of corals. ISME J. 2013;7(1):13–27.

21. Ng FFT, Wheeler E, Greig D, Waltzek TB, Gulland F, Breitbart M. Metagenomic identification of a novel anellovirus in Pacific harbor seal (Phoca vitulina richardsii) lung samples and its detection in samples from multiple years. J Gen Virol. 2011;92(6):1318–23. doi: 10.1099/vir.0.029678-0.

22. Hewson I, Johnson MR, Tibbetts IR. An unconventional flavivirus and other RNA viruses in the sea cucumber (Holothuroidea; Echinodermata) virome. Viruses. 2020;12:1057.

23. Jackson EW, Wilhelm RC, Johnson MR, Lutz HL, Danforth I, Gaydos JK, et al. Diversity of sea star-associated densoviruses and transcribed endogenized viral elements of densovirus origin. J Virol. 2020:DOI: 10.1128/JVI.01594-20. doi: 10.1101/2020.08.05.239004.

24. Quast C, Pruesse E, Yilmaz P, Gerken J, Schweer T, Yarza P, et al. The SILVA ribosomal RNA gene database project: improved data processing and web-based tools. Nucleic Acids Res. 2012;41(D1):D590–D6. doi: 10.1093/nar/gks1219.

25. Waldron FM, Stone GN, Obbard DJ. Metagenomic sequencing suggests a diversity of RNA interference-like responses to viruses across multicellular eukaryotes. PLOS Genetics. 2018;14(7):e1007533. doi: 10.1371/journal.pgen.1007533.

26. Kang YJ, Huang W, Zhao AL, Lai DD, Shao L, Shen YQ, et al. Densoviruses in oyster Crassostrea ariakensis. Arch Virol. 2017;162(7):2153-7. Epub 2017/03/28. doi: 10.1007/s00705-017-3343-z. PubMed PMID: 28342032.

27. Richard JC, Leis E, Dunn CD, Agbalog R, Waller D, Knowles S, et al. Mass mortality in freshwater mussels (Actinonaias pectorosa) in the Clinch River, USA, linked to a novel densovirus. Sci Rep. 2020;10(1):14498. doi: 10.1038/s41598-020-71459-z.

28. Guo H, Zhang J, Hu Y. Complete sequence and organization of Periplaneta fuliginosa densovirus genome. Acta Virol. 2000;44(6):315–22.

29. Bochow S, Condon K, Elliman J, Owens L. First complete genome of an Ambidensovirus; Cherax quadricarinatus densovirus, from freshwater crayfish Cherax quadricarinatus. Mar Genomics. 2015;24 Pt 3:305-12. Epub 2015/08/14. doi: 10.1016/j.margen.2015.07.009.

30. Phan TG, Messacar K, Dominguez SR, da Costa AC, Deng X, Delwart E. A new densovirus in cerebrospinal fluid from a case of anti-NMDA-receptor encephalitis. Arch Virol. 2016;161(11):3231-5. Epub 2016/08/16. doi: 10.1007/s00705-016-3002-9.

31. Jackson EW, Pepe-Ranney C, Johnson MR, Distel DL, Hewson I. A highly prevalent and pervasive densovirus discovered among sea stars from the north american Atlantic coast. Appl Environ Microbiol. 2020;86(6). doi: 10.1128/AEM.02723-19.

32. Gudenkauf BM, Eaglesham JB, Aragundi WM, Hewson I. Discovery of urchin-associated densoviruses (Parvoviridae) in coastal waters of the Big Island, Hawaii. J Gen Virol. 2014;95:652–8.

33. Frickhofen N, Young NS. Persistent parvovirus B19 infections in humans. Microb Pathogen. 1989;7(5):319–27. doi: https://doi.org/10.1016/0882-4010(89)90035-1.

34. Molthathong S, Jitrakorn S, Joyjinda Y, Boonchird C, Witchayachamnarnkul B, Pongtippatee P, et al. Persistence of Penaeus stylirostris densovirus delays mortality caused by white spot syndrome virus infection in black tiger shrimp (Penaeus monodon). BMC Vet Res. 2013;9(1):33. doi: 10.1186/1746-6148-9-33.

35. Liu H, Fu Y, Xie J, Cheng J, Ghabrial SA, Li G, et al. Widespread endogenization of densoviruses and parvoviruses in animal and human genomes. J Virol. 2011;85(19):9863–76. doi: 10.1128/jvi.00828-11.

36. Newton AL, Smolowitz R. Chapter 41 - Invertebrates. In: Terio KA, McAloose D, Leger JS, editors. Pathology of Wildlife and Zoo Animals: Academic Press; 2018. p. 1019–52.

37. Mutuel D, Rayallec M, Chabi B, Multeau C, Salmon JM, Fournier P, et al. Pathogenesis of Junonia coenia densovirus in Spodoptera frugiperda: A route of infection that leads to hypoxia. Virology. 2010;403(2):137–44. doi: DOI 10.1016/j.virol.2010.04.003.

38. Roekring S, Smith DR. Induction of apoptosis in densovirus infected Aedes aegypti mosquitoes. J Invert Pathol. 2010;104(3):239–41. doi: DOI 10.1016/j.jip.2010.04.002.

39. Owens L, La Fauce K, Claydon K. The effect of Penaeus merguiensis densovirus on Penaeus merguiensis production in Queensland, Australia. J Fish Dis. 2011;34(7):509–15. doi: DOI 10.1111/j.1365-2761.2011.01263.x.

40. Molthathong S, Jitrakorn S, Joyjinda Y, Boonchird C, Witchayachamnarnkul B, Pongtippatee P, et al. Persistence of Penaeus stylirostris densovirus delays mortality caused by white spot syndrome virus infection in black tiger shrimp (Penaeus monodon). BMC Vet Res. 2013;9. doi: Artn 33 Doi 10.1186/1746-6148-9-33.

41. Nerva L, Forgia M, Ciuffo M, Chitarra W, Chiapello M, Vallino M, et al. The mycovirome of a fungal collection from the sea cucumber Holothuria polii. Virus Res. 2019;273:197737. doi: https://doi.org/10.1016/j.virusres.2019.197737.

42. Wei X, Feng C, Li XH, Mao XX, Luo HB, Zhang DM, et al. Enantiomeric polyketides from the starfish-derived symbiotic fungus Penicillium sp. GGF16-1-2. Chem Biodiv. 2019;16(6). doi: 10.1002/cbdv.201900052.

43. FioRito R, Leander C, Leander B. Characterization of three novel species of Labyrinthulomycota isolated from ochre sea stars (Pisaster ochraceus). Mar Biol. 2016;163(8):10. doi: 10.1007/s00227-016-2944-5.

44. Franco OP, Patino GS, Ortiz AA. Antibacterial and antifungal activity of the starfish Oreaster reticulatus (Valvatida: Oreasteridae) and the sea urchins Mellita quinquiesperforata (Clypeasteroida: Mellitidae) and Diadema antillarum (Diadematoida: Diadematidae) from the Colombian Caribbean. Revista Biol Trop. 2015;63:329–37.

45. Palagiano E, Zollo F, Minale L, Iorizzi M, Bryan P, McClintock J, et al. Isolation of 20 glycosides from the starfish Henricia downeyae, collected in the Gulf of Mexico. J Nat Prod. 1996;59(4):348–54. doi: 10.1021/np9601014.

46. Choi DH, Shin S, Park IK. Characterization of antimicrobial agents extracted from Asterina pectinifera. Int J Antimicrob Agent. 1999;11(1):65–8. doi: 10.1016/s0924-8579(98)00079-x.

47. Chludil HD, Seldes AM, Maier MS. Antifungal steroidal glycosides from the Patagonian starfish Anasterias minuta: Structure-activity correlations. J Nat Prod. 2002;65(2):153–7. doi: 10.1021/np010332x.

48. Mendoza L, Taylor JW, Ajello L. The class Mesomycetozoea: A group of microorganisms at the animal-fungal boundary. Ann Rev Microbiol. 2002;56:315–44. doi: 10.1146/annurev.micro.56.012302.160950.

49. Baker GC, Beebee TJC, Ragan MA. Prototheca richardsi, a pathogen of anuran larvae, is related to a clade of protistan parasites near the animal-fungal divergence. Microbiol. 1999;145:1777–84. doi: 10.1099/13500872-145-7-1777.

50. Fredricks DN, Jolley JA, Lepp PW, Kosek JC, Relman DA. Rhinosporidium seeberi: A human pathogen from a novel group of aquatic protistan parasites. Emerg Infect Dis. 2000;6(3):273–82. doi: 10.3201/eid0603.000307.

51. Arkush KD, Mendoza L, Adkison MA, Hedrick RP. Observations on the life stages of Sphaerothecum destruens n. g., n. sp., a mesomycetozoean fish pathogen formally referred to as the rosette agent. J Euk Microbiol. 2003;50(6):430–8. doi: 10.1111/j.1550-7408.2003.tb00269.x.

52. Blazer VS, Hitt NP, Snyder CD, Snook EL, Adams CR. Dermocystidium sp infection in blue ridge sculpin captured in Maryland. J Aquat Ani Health. 2016;28(3):143–9. doi: 10.1080/08997659.2016.1159622.

53. Arnott SA, Dykova I, Roumillat WA, de Buron I. Pathogenic endoparasites of the spotted seatrout, Cynoscion nebulosus: Patterns of infection in estuaries of South Carolina, USA. Parasitol Res. 2017;116(6):1729–43. doi: 10.1007/s00436-017-5449-3.

54. Lord JC, Hartzer KL, Kambhampati S. A nuptially transmitted ichthyosporean symbiont of Tenebrio molitor (Coleoptera: Tenebrionidae). J Euk Microbiol. 2012;59(3):246–50. doi: 10.1111/j.1550-7408.2012.00617.x.

55. Marshall WL, Celio G, McLaughlin DJ, Berbee ML. Multiple isolations of a culturable, motile ichthyosporean (Mesomycetozoa, Opisthokonta), Creolimax fragrantissima n. gen., n. sp., from marine invertebrate digestive tracts. Protist. 2008;159(3):415–33. doi: https://doi.org/10.1016/j.protis.2008.03.003.

56. Marshall WL, Berbee ML. Facing unknowns: Living cultures (Pirum gemmata gen. nov., sp. nov., and Abeoforma whisleri, gen. nov., sp. nov.) from invertebrate digestive tracts represent an undescribed clade within the unicellular opisthokont lineage Ichthyosporea (Mesomycetozoea). Protist. 2011;162(1):33–57. doi: https://doi.org/10.1016/j.protis.2010.06.002.

57. Glockling SL, Marshall WL, Gleason FH. Phylogenetic interpretations and ecological potentials of the Mesomycetozoea (Ichthyosporea). Fungal Ecol. 2013;6(4):237–47. doi: https://doi.org/10.1016/j.funeco.2013.03.005.

